## Supplementary material for "Genome-resolved year-round dynamics reveal a broad range of giant virus microdiversity": Fig. S1 - Fig. S10

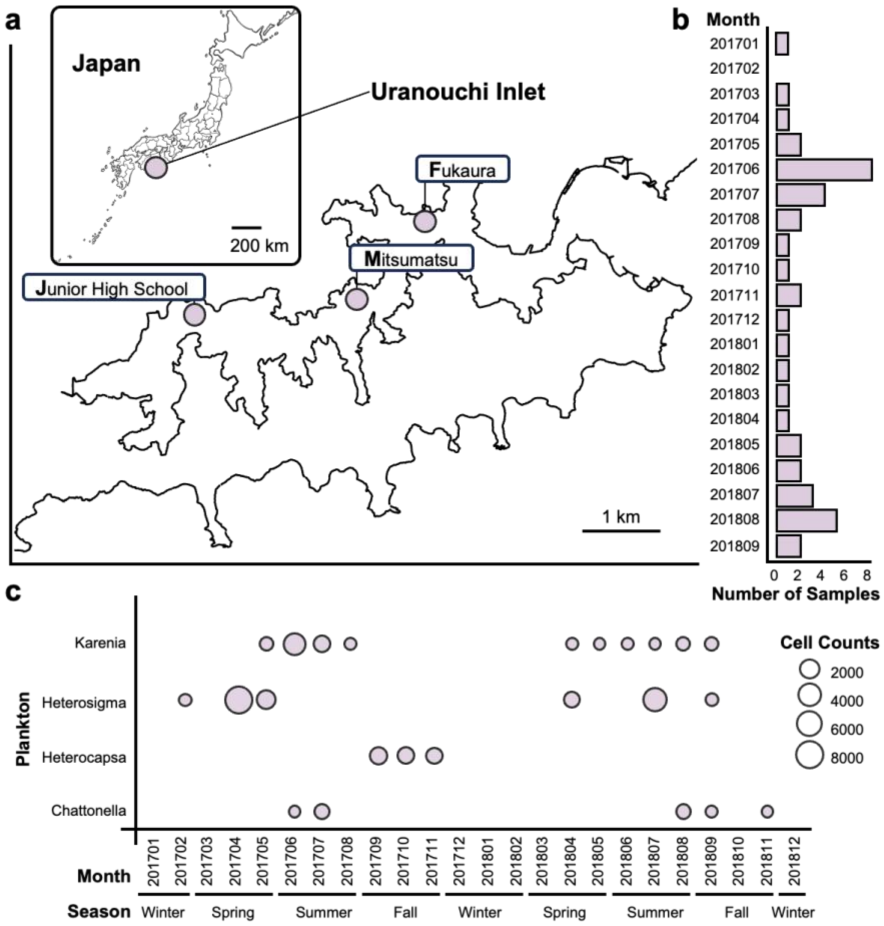


**Figure S1.** Locations of three sampling sites in Uranouchi Inlet, Kochi Prefecture, Japan. Histogram plot in the right panel stands for the sampling frequency in a monthly order.

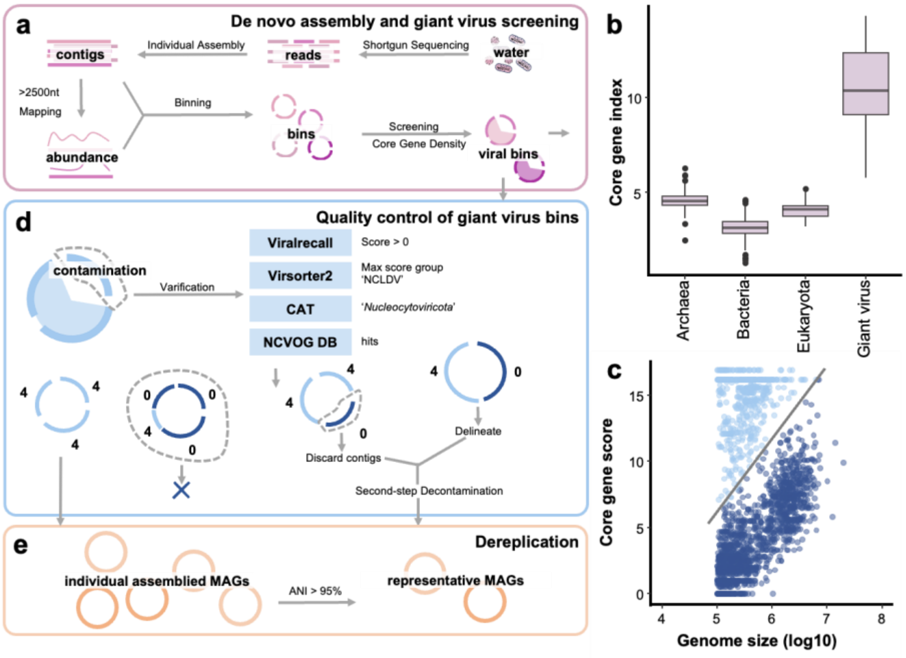


**Figure S2.** (A) Schematic flowchart for generating high-quality MAGs of giant viruses. (B) The distribution of the core gene density index across different groups of organisms and giant viruses (*Nucleocytoviricota*) reference genomes. The threshold for screening giant viruses was determined as 5.75. (C) The distribution of core gene density index of all reference genomes and bins generated in the Uranouchi Inlet. A similar gap was observed (Index = 5.75).

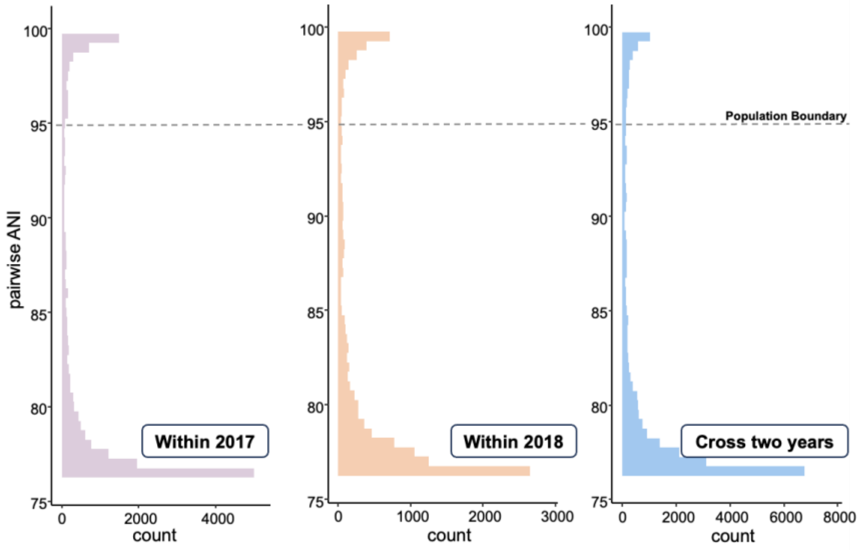


**Figure S3.** Genetic discontinuity observed in the MAGs generated in the Uranouchi Inlet. Histogram plot showing the distribution of ANI values among the MAGs. Only ANI values in the 76–100% range are shown. (A) MAG pairs both originating from the 2017 samples. (B) MAG pairs both originating from the 2018 samples. (C) MAG pairs that originated from the 2017 and 2018 samples, respectively.

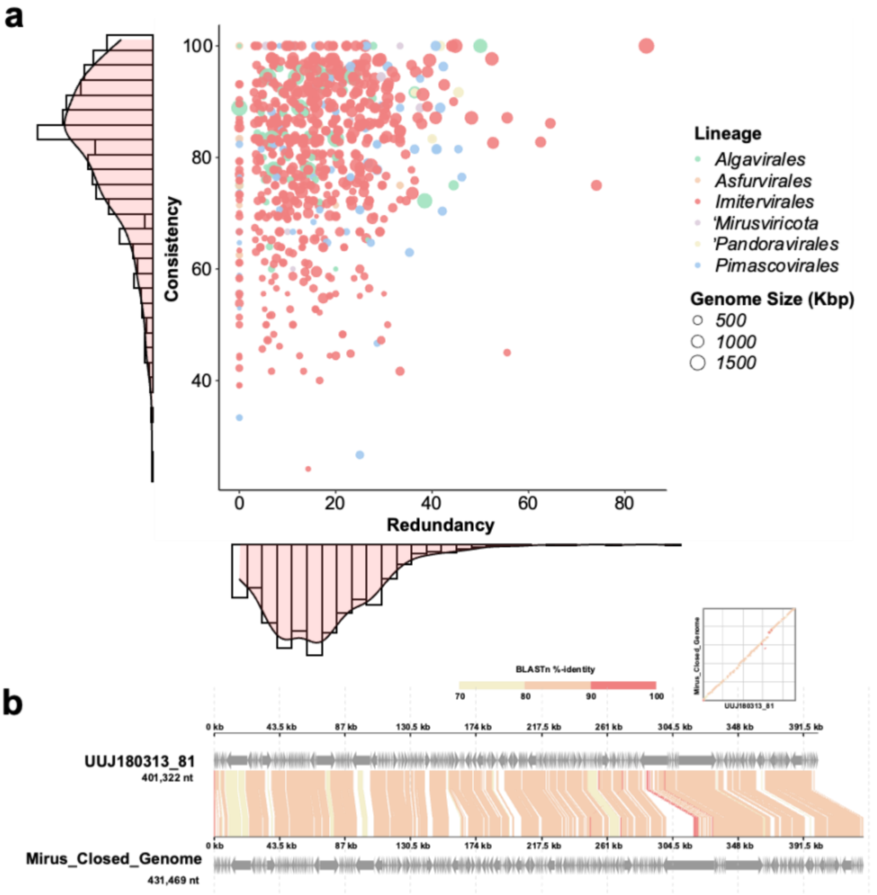


**Figure S4.** (**A**) Distribution of the consistency and redundancy of giant virus MAGs. Color stands for the lineage of giant viruses and scatter size stands for the genome size of each MAG. Two histogram plots demonstrate the distribution of MAG qualities. Medians of the consistency and redundancy were 83.33% and 14.81%, respectively. (**B**) A near complete mirusvirus was assembled from Uranouchi Inlet, demonstrating a considerable genome similarity with the mirusvirus identified in Mediterranean Sea. Contigs of UUJ180313_81 were reordered and concatenated by ReCCO for better visualization. Alignment and visualization were done by DiGAlign.

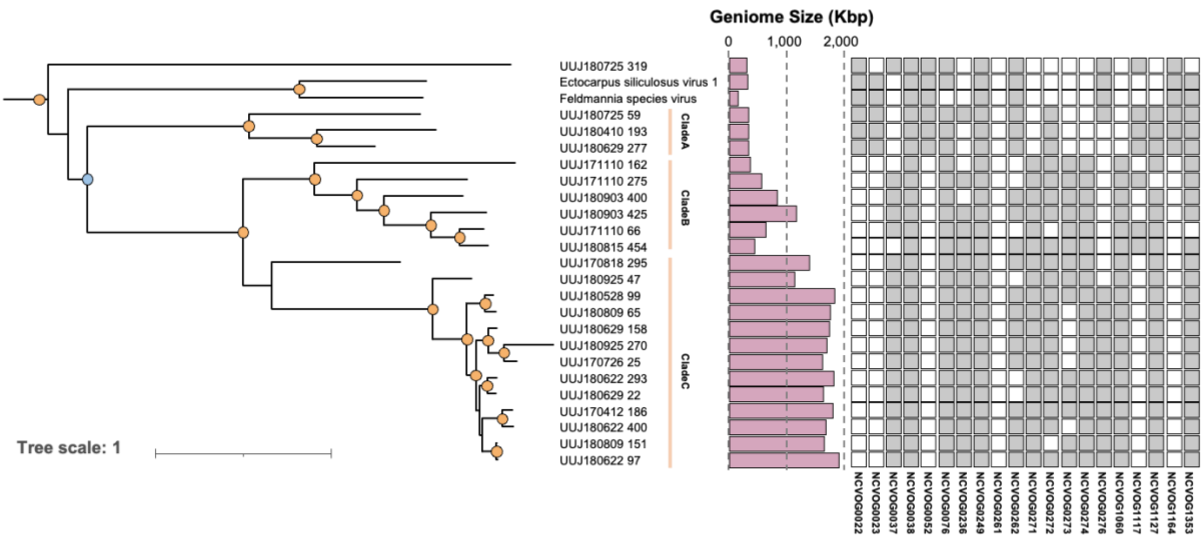


**Figure S5.** A subtree of detected MAGs with a large genome size that are related to EsV and FsV. The presence (grey) and absence (white) of 20 universal genes of giant viruses are shown (ones conserved in novel clades are marked in bold): (1) NCLDV major capsid protein (NCVOG0022);(2) **A32-like packaging ATPase**

(NCVOG0249);(3) Erv1/Alr family disulfide (thiol) oxidoreductase (NCVOG0052);(4) **family B DNA polymerase** (NCVOG0038);(5) D5-like helicase-primase (NCVOG0023);(6) **DNA topoisomerase II** (NCVOG0037);(7) **FLAP-like endonuclease XPG** (NCVOG1060);(8) **DNA or RNA helicases of superfamily II** (NCVOG0076);(9) **Poxvirus late transcription factor VLTF3** (NCVOG0262);(10) Poxvirus late transcription factor VLTF2 (NCVOG1164);(11) **Poxvirus early transcription factor VETF** (NCVOG0261);(12) **Transcription initiation factor IIB** (NCVOG1127);(13) **Transcription factor S-II** (TFIIS) (NCVOG0272);(14) **DNA-directed RNA polymerase subunit alpha** (NCVOG0274);(15) **DNA-directed RNA polymerase subunit beta** (NCVOG0271);(16) **DNA-directed RNA polymerase subunit 5** (NCVOG0273);(17) mRNA capping enzyme, guanylyltransferase (NCVOG1117);(18) mRNA capping enzyme, methyltransferase (NCVOG1117);(19) **Nudix hydrolase** (NCVOG0236);(20) **ribonucleoside diphosphate reductase, large subunit** (NCVOG1353);(21) **ribonucleoside diphosphate reductase, beta subunit** (NCVOG0276). Nodes with UFbootstrap values >= 95 and >= 85 are marked with yellow and blue circles, respectively.

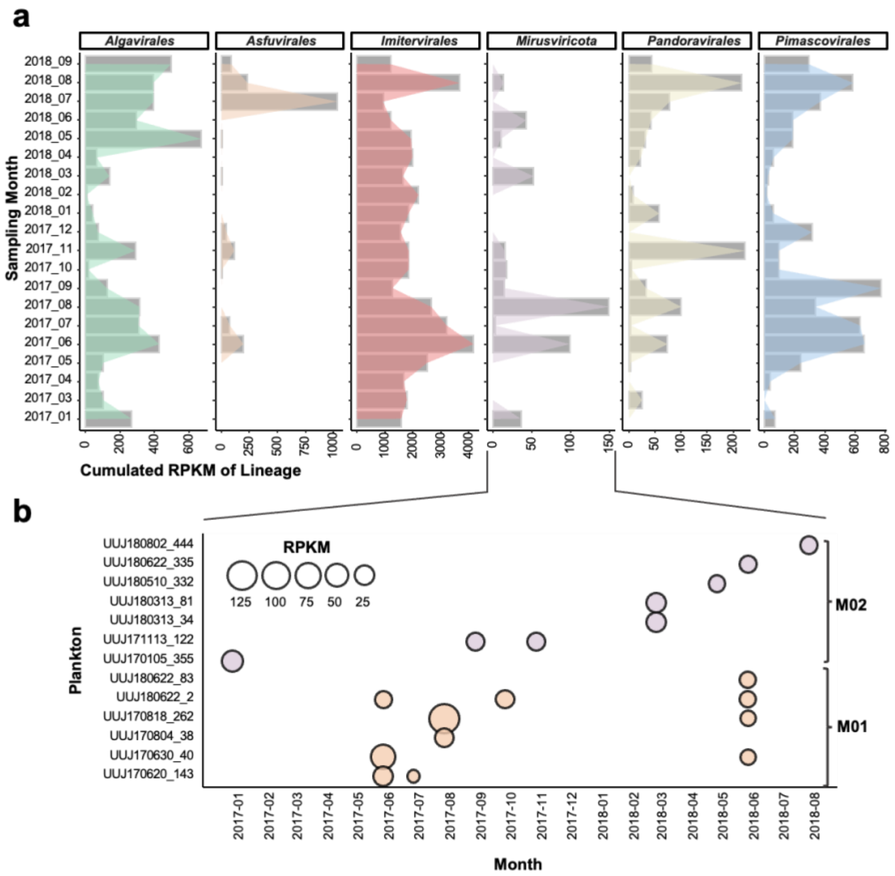


**Figure S6.** (**A**) Cumulative abundance dynamics of GV lineages from January 2017 to September 2018. The cumulative abundance of each GV lineage was calculated by summing up the RPKM of all the MAGs belonging to the lineage. (**B**) RPKM dynamics of *Mirusviricota* from January 2017 to September 2018. RPKM of MAGs in the given samples were given by CoverM based on reads mapping.

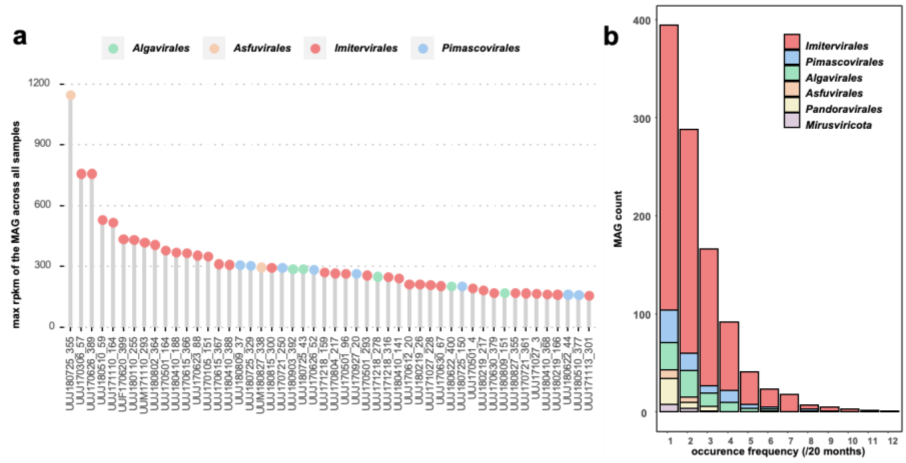


**Figure S7.** Ecological characteristics of individual MAGs. (**A**) Top 50 giant viruses exhibiting the highest maximum RPKM (representing their peak abundance in one sample) are showcased. The x-axis labels represent the MAG IDs, with colors indicating the lineage of each MAG. (**B**) The distribution of giant virus populations across various occurrence frequencies is presented, calculated over a 20-month period from January 2017 to September 2018.

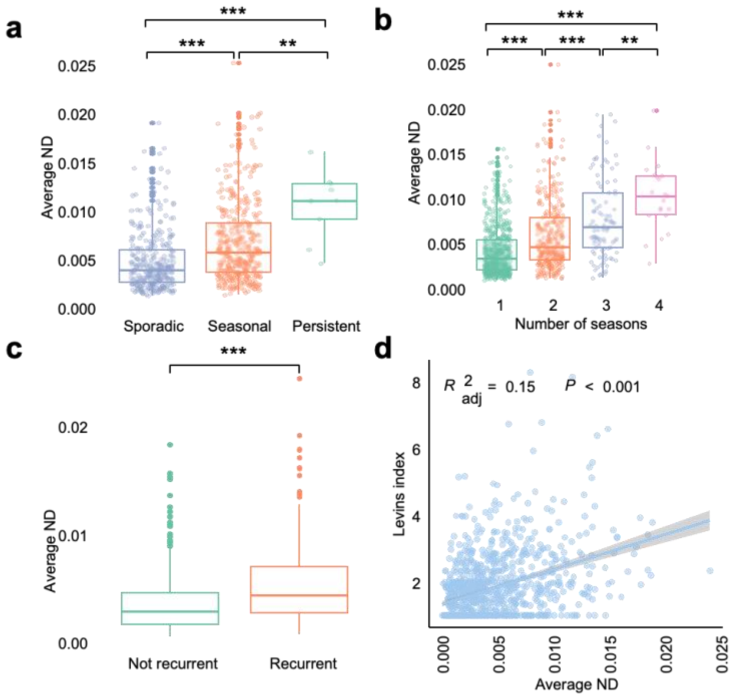


**Figure S8**. Temporal dynamics and microdiversity of giant viruses. (**A**) Average nucleotide diversity of viruses in three niche categories: Persistent, Seasonal, and Sporadic. (**B**) Average nucleotide diversity of viruses that occurred in one, two, three, or all four seasons. (**C**) Comparison of average nucleotide diversity of viruses that only occurred in 2017 versus those that occurred in both 2017 and 2018. P values for (**A**), (**B**), and (**C**) at the top were obtained using the Kruskal-Wallis rank sum test. P values < 0.01 are shown as **, and P values < 0.001 are shown as ***. (**D**) Correlation between niche breadth (Levins index) and the average nucleotide diversity of all giant virus MAGs. The values of nucleotide diversity were determined using InStrain v1.0.0. The adjusted R-squared and p value were provided by the 'stat_poly_eq()' function in the 'ggpmisc' package in R.

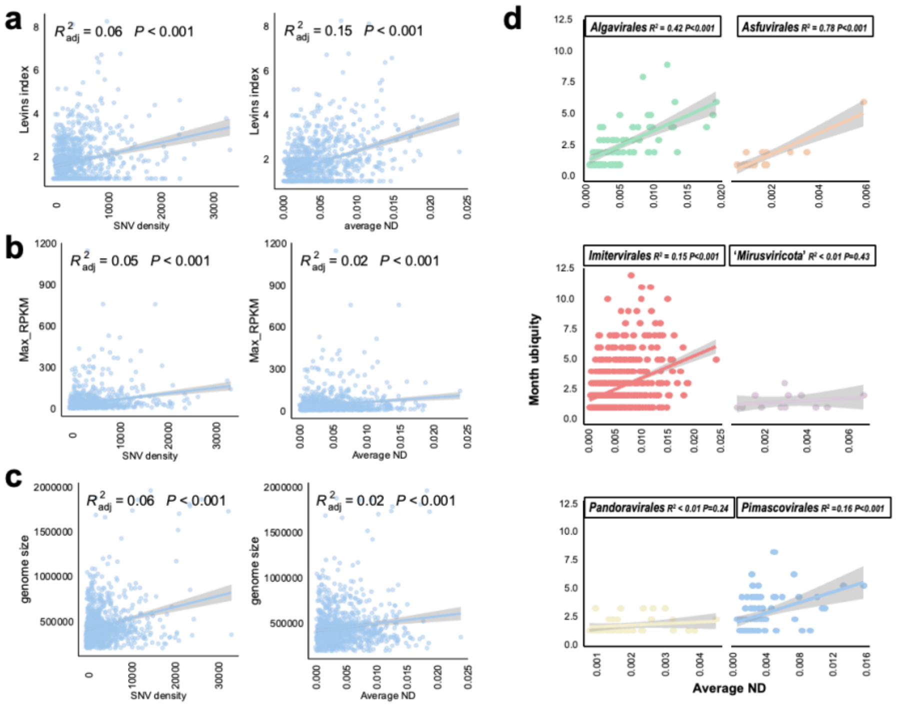


**Figure S9.** The correlation between SNV density and (**A**) Levins index, (**B**) Maximum RPKM and (**C**) Genome size. (**D**) Levins index vs. average nucleotide diversity of giant virus MAGs for each lineage. The adjusted R-squared and p value were provided by the 'stat_poly_eq()' function in the 'ggpmisc' package in R.

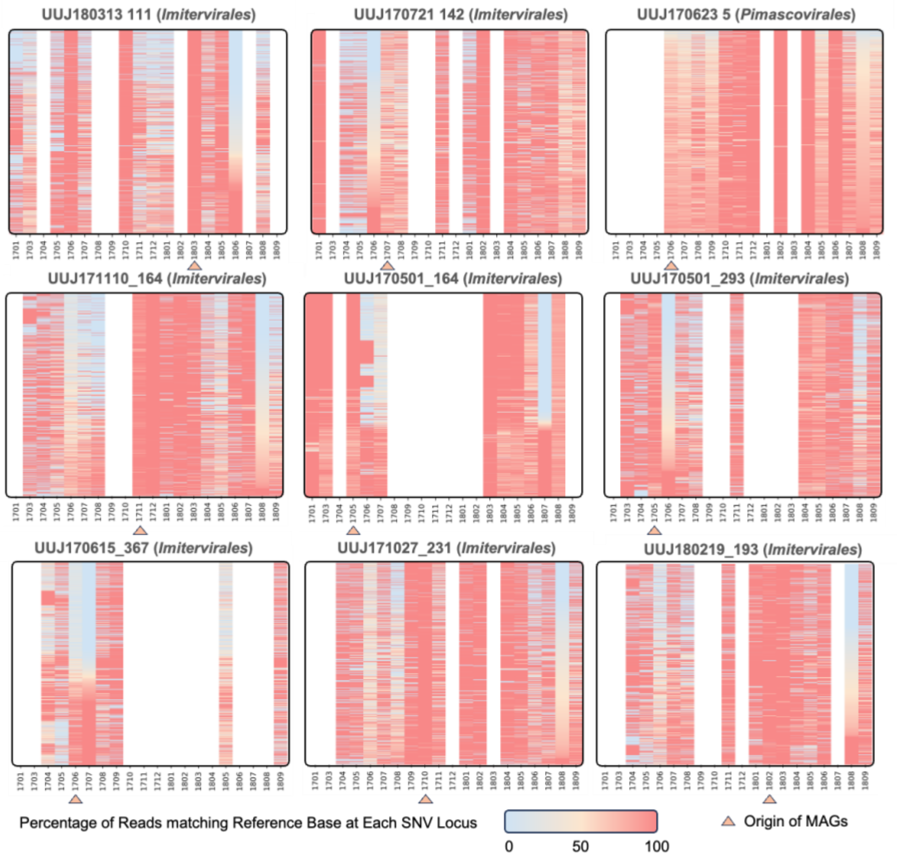


**Figure S10.** Temporal dynamics of SNV allele frequencies within 9 "Persistent" populations. SNV positions are arrayed along the y axis, with each row representing one SNV site. Color indicates allele frequency, that is, the percentage of metagenomic reads supporting the reference allele during each time period. Within a given MAG, if an SNV site is observed in more than 30% of the months during which the MAG has an RPKM greater than 0 covering more than 50% genome length, the SNV site is retained for allele frequency calculation and shown here. White represents an incomparable SNV (either there are not enough reads mapping or the SNV site is not prevalent across temporal samples). The reference bases are derived from the MAG sequence, and the sites are sorted by the month with the minimum average similarity to the reference. Triangle represents origin of the MAG.
