## Supplementary information for "Genome-resolved year-round dynamics reveal a broad range of giant virus microdiversity"

#### Introduction

In this study, we designed an automatic pipeline to generate metagenome-assembled genomes (MAGs) of giant viruses. The pipeline includes the following steps: 1) individual metagenome assembly and binning, 2) screening for giant viruses, 3) refinement of giant virus MAGs, and 4) deduplication of MAGs (see Methods; Fig. S2). Additionally, we developed the Phylogeny-Informed MAG Assessment (PIMA) to quantitatively evaluate the quality of giant virus MAGs. By combining CheckV[1], a well-accepted tool for estimating the completeness of viral genomes, we assessed the quality of the assembled MAGs (Fig. S4; Table S3). However, the quality of viral genomes can be influenced by several factors, like sampling approach, experimental methods, and sequencing depth. Assessing the effectiveness of the pipeline itself is challenging. Different pipelines can produce varying results and interpretations from the same data. Therefore, an additional evaluation is necessary to provide a comprehensive assessment of the performance of the pipeline used in this study, comparing it against established pipelines and ensuring its robustness in recovering giant virus lineages.

In the evaluation work, by leveraging 28 *Tara* Arctic metagenomes of protist size fractions, we compared our pipeline against two established pipelines used in global giant virus metagenomic sruveys[2,3]. We first assessed the completeness and contamination of MAGs generated by the three pipelines using CheckV. Next, by aligning MAGs from the three pipelines with a manually curated virus genome dataset, we investigated the ability of each pipeline to recover viral lineages, including the novel virus phylum, *Mirusviricota*. Our benchmark results suggested that the pipeline used in this study offers a balanced and fine approach, capable of reconstructing clean genomes and recovering diverse viral lineages.

### Method

#### Data

To evaluate the performance of the binning pipeline used in this study, we analyzed a *Tara* Arctic metagenome dataset to benchmark our pipeline against two earlier metagenomic surveys. This dataset was used in a previous project including a manual curation step[4]. This dataset comprised 28 metagenomes of 17 *Tara* Oceans stations with filter sizes larger than 0.8 µm. The sequencing data and metadata are available in the European Nucleotide Archive (ENA) database under Study Accession PRJEB9691. The Run Accessions are as follows: ERR4691596, ERR4691597, ERR4691603, ERR4691606, ERR4691607, ERR4691614, ERR4691615, ERR4691621, ERR4691622, ERR4691627, ERR4691630, ERR4691632, ERR4691638, ERR4691639, ERR4691646, ERR4691647, ERR4691657, ERR4691661, ERR4691662, ERR4691668, ERR4691673, ERR4691674, ERR4691681, ERR4691685, ERR4691688, ERR4691690, ERR4691692, ERR4691694.

#### Generation of Giant Virus Genomes using different pipelines

Three pipelines were employed to generate the MAGs of selected metagenomes. Before binning, contigs were assembled using MEGAHIT v1.2.9[5] with a k-mer range from 31 to 141, with a step of 10. Bowtie2 v2.4.5[6] mapped reads to assembled contigs longer than 1 Kbp, which were then sorted and converted to BAM files using samtools v1.16[7]. The pipeline used in this study (for generating giant virus MAGs from the Uranouchi Inlet) is described in the Methods section of the main text. For the other two pipelines, we followed the methods documented in the literature to replicate their binning approaches as closely as possible.

The first pipeline[2], referred to as Pipeline1, removed outliers in the phylogenetic tree through manual inspection. To remove outliers, we used a tetranucleotide frequency-based method, excluding outliers from each lineage (those with values $<-2.5$ or $>2.5$ s.d. within each genome bin), following another previous work[8]. The other pipeline, referred to as Pipeline2, provided detailed parameters in the publication[3]. We adhered to the software and parameters used for both initial binning and screening steps as previously described.

#### Benchmark

Completeness and contamination were evaluated using CheckV v1.0.1[1]. Additionally, we aligned MAGs from the three pipelines with a manually curated GOEV virus genome dataset[4]. The genome-wise alignment was performed using fastANI v1.33 with an ANI threshold of 95%. The taxonomy was determined by the aligned MAGs in the GOEV database. Gene calling was done using Prodigal v2.6.3[9], and ortholog groups (OGs) were identified using OrthoFinder v2.5.5[10].

#### Result and Discussion

A total of 498 MAGs were generated by the three pipelines. The pipeline used in this study produced 153 MAGs, while the other two pipelines generated 312 and 33 MAGs, respectively (Fig. SI1). All three pipelines produced genomes with genome sizes around 300 Kbp. Notably, the pipeline from this study yielded MAGs with the highest median genome size (~357 Kbp). Meanwhile, the largest genome, with a size of 2,417 Kbp, was generated by Pipeline1.

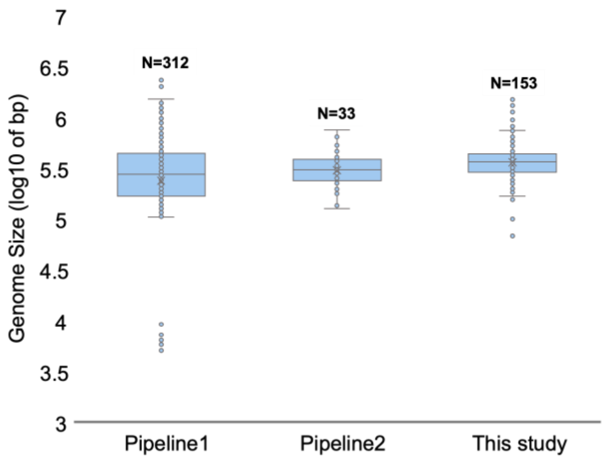


**Figure SI1.** Genome size of MAGs generated using three pipelines.

We then used CheckV to assess the quality of the yielded MAGs. Three algorithms for calculating the completeness (AAI, HMM_upper, HMM_lower) provided a similar ranking among three pipelines. Pipeline1 had a higher median completeness than the pipeline used in this study and Pipeline 2 (Fig. SI2a, b, c). However, the distribution range of completeness by Pipeline1 was also broader than that of the other two pipelines, suggesting there are many low complete MAGs. Additionally, the estimated contamination of Pipeline1 was higher than that of the other two pipelines (Fig. SI2d). Pipeline2, described as stringent, demonstrated a notably low contamination rate.

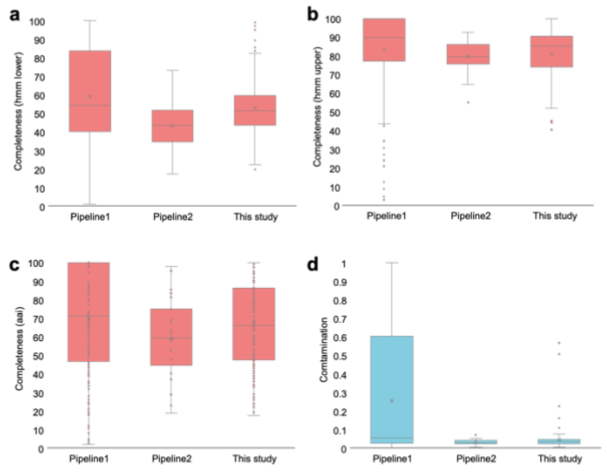


**Figure SI2.** Quality assessment of MAGs generated using three pipelines. Completeness and contamination were assessed using CheckV. Three algorithms for calculating the completeness (**A**) HMM-based (lower); (**B**) HMM-based (upper); (**C**) AAI-based. (**D**) Estimated contamination rate of MAGs.

Both completeness and contamination results suggested the pipeline used in this study produced MAGs of moderate quality. Considering the number of MAGs, pipeline of this study was able to control the contamination rate while generating a sufficient number of MAGs, making it an ideal choice in giant virus study.

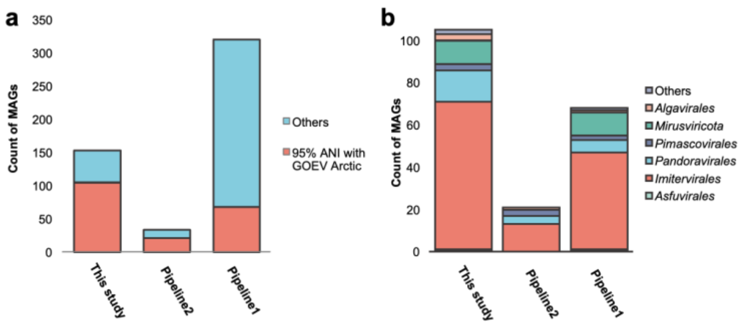


**Figure SI3.** Comparison of MAGs yielded in this benchmark with the GOEV database. (**A**) Number of MAGs that share over 95% ANI with MAGs in the GOEV database. (**B**) Number of MAGs of each lineage among the MAGs aligned to the GOEV database.

We then aligned the MAGs yielded by the three pipelines with those from the GOEV database. Since the metagenomic sequences used for binning were included in the GOEV database, the MAGs in the benchmark were expected to be represented in the GOEV database. The alignment was performed using fastANI with an ANI threshold of 95%, which is considered as the boundary of species level[11,12].

Notably, while the pipeline used in this study did not generate the largest number of MAGs (Fig. SI3a), it did produce the highest number of MAGs found in the GOEV database (N = 105; 68.62%). Pipeline2 also recovered a high ratio of GOEV Arctic MAGs (63.63%). However, only 21.25% of the MAGs yielded by Pipeline1 were represented in the GOEV database.

Out of six major lineages, *Algavirales*, *Pimascovirales*, *Pandoravirales*, and *Imitervirales* were detected by all three pipelines. *Mirusviricota* and *Asfuvirales* were only detected by the pipeline used in this study and Pipeline1 (Fig. SI3b).

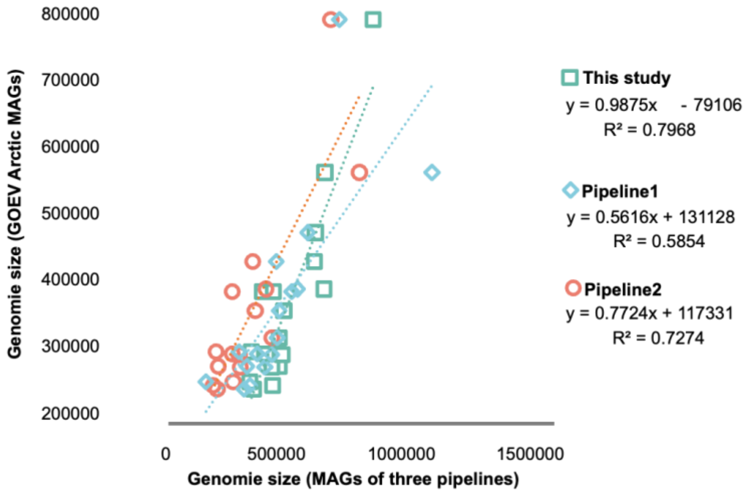


**Figure SI4.** Comparison of genome sizes of MAGs yielded by the three pipelines with the GOEV database.

We then checked the genome consistency, assuming that the GOEV database provided reasonable genome sizes due to manual curation and delineation. Sixteen groups of quadruple genome consensuses were detected (i.e., all consensuses included MAGs from the three pipelines and one GOEV MAG). We compared the genome sizes and found that the pipeline used in this study showed better consistency with GOEV Arctic MAGs, with a slope close to one (0.9875) and the highest R-squared value (0.7968).

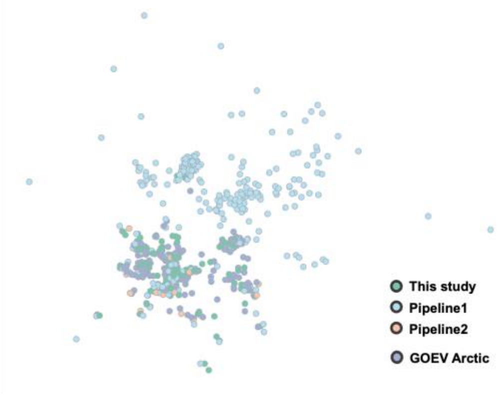

**Figure SI5.** Gene-sharing patterns of the MAGs from the three pipelines and MAGs in the GOEV database. Each node represents a MAG, and nodes are connected by shared orthologous groups (OGs).

Most of the MAGs generated by Pipeline1 shared distant functional clusters with other MAGs (Fig. SI5). Meanwhile, GOEV MAGs, as well as MAGs from Pipeline2 and this study, shared a similar gene repertoire. These results, along with the ANI alignment (Fig. SI3), indicate that Pipeline1 demonstrated the potential for discovering new viral species and functions. However, the GOEV database, which employs a phylogeny-guided single-gene search to identify giant viruses, also exhibited strong capabilities in uncovering new viruses[4]. Therefore, our pipeline possesses advantages comparable to other methods.

Overall, the pipeline used in this study showed a moderate quality compared to the two established pipelines. One pipeline was very stringent, representing the most conserved parts of giant virus genomes[3]. Meanwhile, the other one demonstrated potential by recovering novel virus genomes[2]. Moreover, the pipeline used in this study could recover all major lineages, including the new phylum, *Mirusviricota*, and showed the highest genome size consistency of MAGs with the GOEV database. Therefore, we believe the pipeline of this study offers a good trade-off, balancing quality and high sensitivity in detecting giant virus genomes. The MAGs generated from Uranouchi also suggest that MAGs produced using this pipeline have good quality, making it suitable for generating representatives for microdiversity analyses.
